# The consequence of ATP synthase dimer angle on mitochondrial morphology studied by cryo-electron tomography

**DOI:** 10.1101/2023.02.02.526626

**Authors:** Emma Buzzard, Mathew McLaren, Piotr Bragoszewski, Andrea Brancaccio, Holly Ford, Bertram Daum, Patricia Kuwabara, Ian Collinson, Vicki A.M. Gold

**Affiliations:** Living Systems Institute, University of Exeter, Exeter, UK; Faculty of Health and Life Sciences, University of Exeter, Exeter, UK; Nencki Institute of Experimental Biology, Polish Academy of Sciences, Warsaw, Poland; Institute of Chemical Sciences and Technologies “Giulio Natta”, Department of Chemical Sciences and Materials Technologies, National Research Council (CNR), Rome, Italy; School of Biochemistry, University of Bristol, BS8 1TD, UK

**Author notes:** corresponding author: Vicki Gold.

**Keywords:** Mitochondria, ATP synthase, Cryo-electron tomography, Sub-tomogram averaging, AlphaFold

## Abstract

Mitochondrial ATP synthases form rows of dimers, which induce membrane curvature to give cristae their characteristic lamellar or tubular morphology. The angle formed between the central stalks of ATP synthase dimers varies between species. Using cryo-electron tomography and sub-tomogram averaging, we determined the structure of the ATP synthase dimer from the nematode worm *C. elegans* and show that the dimer angle differs from previously determined structures. The consequences of this species-specific difference at the dimer interface were investigated by comparing *C. elegans* and *S. cerevisiae* mitochondrial morphology. We reveal that *C. elegans* has a larger ATP synthase dimer angle with more lamellar (flatter) cristae when compared to yeast. The underlying cause of this difference was investigated by generating an atomic model of the *C. elegans* ATP synthase dimer by homology modelling. A comparison of our *C. elegans* model to an existing *S. cerevisiae* structure reveals the presence of extensions and rearrangements in *C. elegans* subunits associated with maintaining the dimer interface. We speculate that increasing dimer angles could provide an advantage for species that inhabit variable-oxygen environments by forming flatter more energetically efficient cristae.

## Main Text

### Introduction

The F_1_F_o_ ATP synthase is a molecular motor ubiquitous to all living organisms, required for the essential conversion of an electrochemical gradient into the universal energy currency ATP (1). The ATP synthase is composed of a catalytic F_1_ head connected to a membrane-embedded F_o_ motor by a central stalk; the entire assembly is visualised as a lollipop shape when examined by electron microscopy (2,3). The central stalk transmits the torque generated by rotation of F_o_ to the F_1_ head, and a peripheral stalk acts as an elastic spring, ensuring malleable coupling between F_1_ and F_o_ (4). Mitochondrial ATP synthases across species share the same complement of core subunits with varying nomenclature (Table S1) (5,6). In metazoans studied to date, the F_1_ head is comprised of α and β subunits, the central stalk of γ, δ and ε subunits, the peripheral stalk of b d, F_6_ and oligomycin sensitivity conferral protein (OSCP) subunits, and the F_o_ motor contains the c-ring and subunit a.

Mitochondrial ATP synthases can assemble into dimers (7), of which there are 4 types (8): Type I is present in both multicellular (9–11) and unicellular organisms (12) and types II-IV are present in various unicellular organisms (13–18), reviewed in (8). When compared to type II-IV dimers, previously studied type I dimers contain an additional set of subunits at the dimer interface: e, f, g, i/j, k and 8 (Table S1) (8). Based on biochemical and imaging experiments, subunits e and g were shown to be essential for dimer formation (7,11,19–21). Dimers of ATP synthase assemble into oligomeric rows (or ribbons) along the curved ridges of crista membranes, observed by cryo-electron tomography (cryoET) (9,11,22). This formation of dimer rows is mediated by an ancestral motif in subunits e and g (20,21) with assistance from subunit k (5,23,24). Formation of dimer rows is required for crista membrane curvature, and thus maintenance of lamellar or tubular shaped cristae (11,12). Deformation of cristae into balloon-like structures was observed in *S. cerevisiae* after knockdown of interface subunits e or g (11) and in ageing *P. anserina,* when dimers disassociated into monomers (12). Moreover, molecular simulations indicated that ATP synthase dimers have an innate propensity to induce membrane curvature (25). This was confirmed experimentally when dimers reconstituted into liposomes spontaneously self-assembled into oligomeric rows to engender this curvature, maintaining identical dimer angles to those observed in whole mitochondria (26).

*In situ* structures of type I ATP synthase dimers have been determined from native membranes (10,12,25–27). Mammals and fungi both display an average angle between the dimer heads of ∼86° (10). Interestingly, higher-resolution single particle analysis of the purified bovine ATP synthase dimer reveals that dimer angles likely vary around this average (between 76° and 95°), depending on catalytic state (28). Further atomic-detail structures of purified mitochondrial type I ATP synthase dimers have also been determined from mammals (*Bos taurus* (23) and fungi (*S. cerevisiae* (29) and *Y. lipolytica* (22)). Whilst the structure and organisation of ATP synthase dimers has been studied across a range of different species, our knowledge of ATP synthases in invertebrates is lacking. The free-living nematode worm *C. elegans* is a well-established model system for the study of invertebrate cell and developmental biology (30), including the role of mitochondria in metabolism, health, disease and aging (31). To complement *in vivo* physiological studies, intact mitochondria can be stably prepared (32,33) for biochemical and structural analyses (34). Interestingly, studies have shown that nematodes lack the dimer-specific subunits i/j, k (35) and 8 (36) found in mammals and fungi (Table S1). Subunit 8 is encoded by one of two overlapping ATP synthase genes on the mitochondrial genome (37). Proteins encoded on the mitochondrial genome are translated from essential genes (38,39); thus it follows that subunit 8 is likely to be essential for respiration in mammals and fungi. The lack of dimer-specific subunits in *C. elegans* provides a unique opportunity to investigate how certain subunits influence ATP synthase dimer angles and mitochondrial morphology.

In this study, we employ cryoET and sub-tomogram averaging to determine the structure and membrane organisation of the *C. elegans* ATP synthase, revealing a novel average dimer angle of 105°. We also discover extra mass at the dimer interface compared to an equivalent *S. cerevisiae* structure determined in the membrane (11). We subsequently compare mitochondria from both *C. elegans* and *S. cerevisiae* to investigate the relationship between ATP synthase dimer angle and crista morphology. Finally, we use AlphaFold (40) and AlphaFold multimer (41) to predict how protein chains in the *C. elegans* ATP synthase dimer may be arranged. This allows us to analyse subunit differences at the dimer interface and postulate the cause of variations in angle. We speculate that an evolutionary divergence at the dimer interface and corresponding widening of the dimer angle may be an adaptation to more variable oxygen environments.

## Results

### The architecture of the *C. elegans* ATP synthase dimer

To determine the arrangement and architecture of ATP synthase dimers in *C. elegans*, tomograms of whole mitochondria (Fig. 1A) and of isolated crista membranes (Fig. 1B) were analysed. ATP synthases were unambiguously identified by the characteristic lollipop shape of the 10 nm diameter F_1_ heads positioned ∼10 nm away from the membrane. We confirmed the presence of oligomeric ATP synthase dimer ribbons, localised at the sharp curved ridges of crista membranes, in both samples (Fig. 1A, B). Due to the obscuring presence of a dense matrix in whole mitochondria, many more dimers could be visualised in isolated crista membrane samples. Therefore, 3,234 dimer pairs were extracted from the crista membrane data for sub-tomogram averaging. After classification, a map of the *C. elegans* ATP synthase dimer was determined from 1,755 dimer pairs (Fig. 1C, Fig. S1, S2). Both the central and peripheral stalks were resolved clearly.

**Figure 1.**
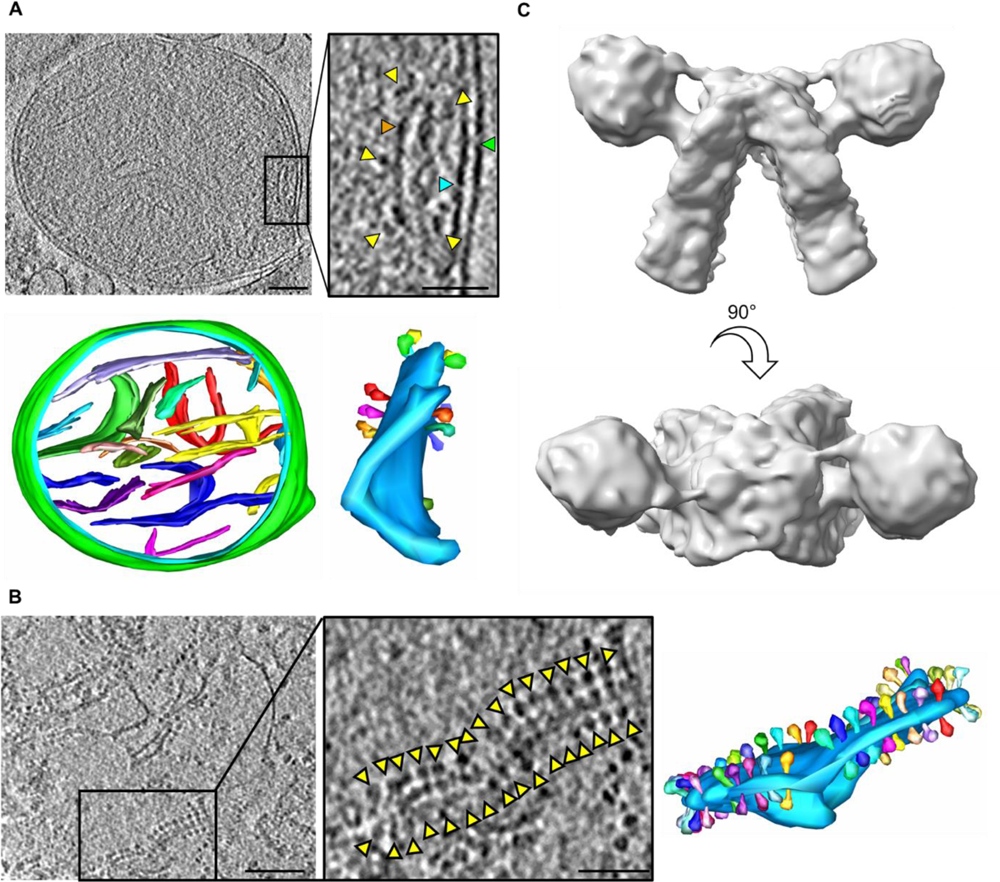
ATP synthase dimer rows, and sub-tomogram average of the ATP synthase dimer from *C. elegans*. **(A)** Tomographic slice through a whole *C. elegans* mitochondrion (top) and corresponding segmentation (bottom; outer membrane green, inner membrane light blue, and a different colour for each crista membrane). The boxed region shows an enlarged image of a single crista membrane, with green, blue and orange arrowheads indicating the outer, inner and crista membranes respectively, and yellow arrowheads indicating ATP synthase F_1_ heads. The crista membrane is coloured light blue in the corresponding segmentation; each ATP synthase dimer pair is coloured differently. **(B)** Tomographic slice through *C. elegans* isolated crista membranes (left, yellow arrowheads indicating ATP synthase F_1_ heads) and corresponding segmentation (right). The boxed region shows an enlarged image of a single crista membrane, with the corresponding segmentation coloured as in panel A. Scale bars, 100 nm for tomograms, and 50 nm for enlarged views of crista membranes. **(C)** Sub-tomogram average of the *C. elegans* ATP synthase dimer. Upper panel shows side view, lower panel shows top-down view.

Previous studies revealed a type I dimer angle of ∼86° across a range of mammalian and fungal species (10,12,25–27). The architecture of the membrane-bound *C. elegans* ATP synthase dimer is unlike any other species studied so far, with an average angle of 105° between the dimer heads (Fig. 2A). A comparison to the structure of the membrane-bound *S. cerevisiae* dimer (42), revealed that the wider dimer angle in *C. elegans* corresponds with a sharper angle of membrane curvature (50° compared to 74°) (Fig. 2B). Accordingly, a shorter distance is measured between the ATP synthase central stalks in *C. elegans* compared to *S. cerevisiae* (16.5 nm compared to 20 nm), which would have the effect of bringing the crista membranes closer together. Intriguingly, the dimer interface in the *C. elegans* map is also visually different to its *S. cerevisiae* counterpart (Fig. 2B), and indeed all other type I dimers studied to date (10,12,25–27). This difference is likely attributable to the different complement of dimer interface subunits present in *C. elegans* compared to *S. cerevisiae* (Table S1, Fig. 2C). We also analysed the inter-dimer distance and angle between dimer heads in consecutive dimers in the oligomeric rows. This revealed an inter-dimer distance of 12.5 nm and angle between dimer heads of 20°. Despite differences in dimer angle, these values are consistent with those reported previously for the type II dimer from green algae (*Polytomella* sp.) (26) (Fig. S3), suggesting that dimer angle does not influence oligomerisation of ATP synthases into rows.

**Figure 2.**
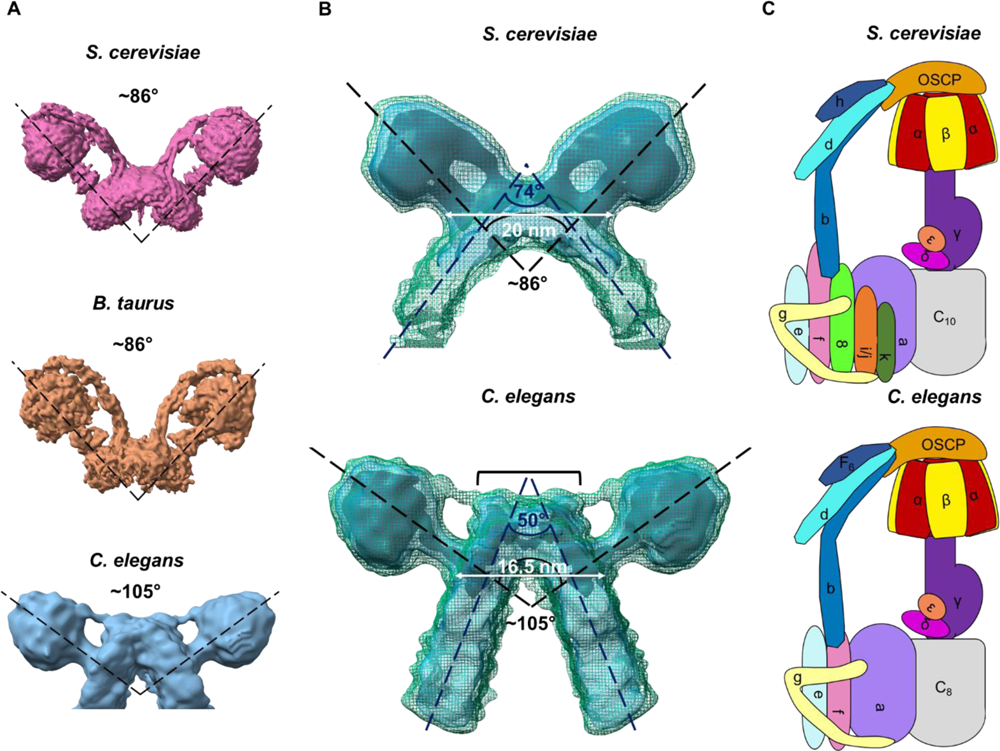
The *C. elegans* ATP synthase compared to other species. **(A)** Structures depicting the range of average dimer angles observed in *S. cerevisiae* [EMD-7067] (29), bovine heart [EMD-11436] (28), and *C. elegans* (this work, [EMD-XXX]), using the highest resolution structures available. **(B)** Direct comparison between *S. cerevisiae* [EMD-2161] (11) and *C. elegans* ATP synthase sub-tomogram averages, with the angle between F_1_ dimer heads, the angle of crista membrane curvature, and distance between the central stalks for each monomer indicated. A bracket highlights the extra mass at the *C. elegans* dimer interface not apparent in *S. cerevisiae*. Black, transparent blue and dark green mesh represent decreasing threshold levels for the averages. **(C)** Cartoon detailing occurrence of ATP synthase subunits in *S. cerevisiae* and *C. elegans*, each labelled with corresponding nomenclature for the species (details in Table S1).

### A wider dimer angle in *C. elegans* corresponds to flatter cristae

We hypothesised that the wider dimer angle associated with sharper membrane curvature in the *C. elegans* ATP synthase dimer (Fig. 2B) would produce flatter cristae with a larger surface area to volume ratio. To test this, tomographic data of whole mitochondria from *C. elegans* and *S. cerevisiae* were collected and quantified. Qualitatively, *C. elegans* mitochondria have more lamellar shaped (or flatter) cristae, with sharp curved ridges, compared to mitochondria from *S. cerevisiae* (Fig. 3A, 3B, Movie S1 & S2). The surface area and volume of the crista membranes were quantified, to reveal that the surface area to volume ratio of the average crista membrane was significantly higher (∼1.5 fold, **** p ≤ 0.0001) in *C. elegans* than in *S. cerevisiae* (Fig. 3B). In accordance with this, the average crista width in *C. elegans* was less than that observed in *S. cerevisiae* (Fig. 3C, D and E), suggesting that dimer angle exerts influence on mitochondrial morphology at the level of membrane curvature.

**Figure 3.**
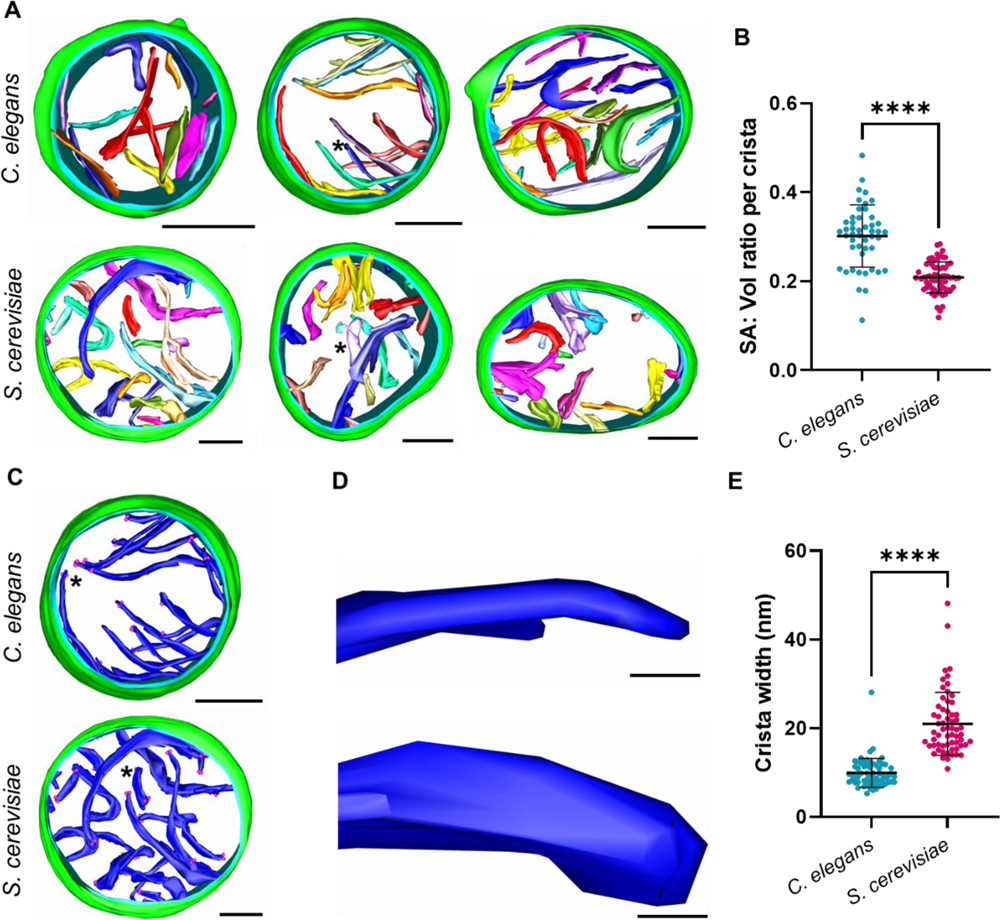
Morphology of mitochondria isolated from *C. elegans* and *S. cerevisiae*. **(A)** Tomographic segmentations of *C. elegans* and *S. cerevisiae* mitochondria are displayed (green, outer mitochondrial membrane; blue, inner mitochondrial membrane; multi-colour, crista membranes). See Movie S1 (*C. elegans*) and Movie S2 (*S. cerevisiae*). **(B)** The mean surface area to volume ratio per crista (n = 3 mitochondria for each organism, with n=47 cristae for *C. elegans* and n=63 cristae for *S. cerevisiae*) was calculated from the segmentations shown in (A). **(C)** A single tomographic segmentation from each organism is shown with all crista coloured blue. Pink dots indicate distances used to measure width. **(D)** Close up of a single crista membrane from each organism (location indicated by asterisks in D to highlight the flatter and thinner crista morphology in *C. elegans* mitochondria compared to *S. cerevisiae*. **(E)** The mean crista width (n= 63 crista tips for *C. elegans* and n= 61 for *S. cerevisiae*) was calculated from the segmentations shown in (A). Error bars in B and E show standard deviation of the mean and significance values were calculated using Welch’s t-test for panel B or using the Mann-Whitney U-test for panel E. **** p ≤ 0.0001. Scale bars in A & C, 200 nm; in D, 20nm.

Mitochondria are dynamic organelles, and crista morphology can be influenced by a wide range of factors such as metabolic state (43–45). However, the average ATP synthase dimer angle remains consistent when imaged in membranes or on purification in detergent (10,28). Nevertheless, we corroborated our findings in whole mitochondria by measuring the width of isolated cristae containing either *C. elegans* or *S. cerevisiae* ATP synthase dimers used for structural determination (Fig. 2B). Our results confirm the narrower crista width in *C. elegans* compared to *S. cerevisiae.* This indicates that the dimer angle and corresponding angle of membrane curvature is consistent, irrespective of the method employed for sample preparation or analysis.

### A unique arrangement of subunits at the *C. elegans* dimer interface

We observed extra mass at the *C. elegans* dimer interface (Fig. 2B) not previously observed in other type I structures determined to date (10). Nematodes are missing subunit 8 (36) (Table S1, Fig. 2C), which plays a key structural role in other species (22,23,29,46). Moreover, subunit 8 is considered essential for respiration (38,39). Therefore, it is likely that other subunits undergo rearrangements at the dimer interface to compensate for the lack of subunit 8 in nematodes, which could contribute to the observed change of dimer angle. To explore this possibility, we performed multisequence alignments with *C. elegans, S. cerevisiae* and *B. taurus* (47–49). This revealed significant extensions in 3 *C. elegans* subunits located at the dimer interface (e, f and g), and in 3 of the 4 subunits in the peripheral stalk (b, d and F_6_) (Table S1, Fig. S4). Mass spectrometry was used to confirm that the extensions identified by sequence in the dimer interface subunits are present in the mature proteins (Fig. S5).

To investigate if the changes in the dimer interface and peripheral stalk subunits could account for the extra mass observed at the dimer interface (Fig. 2B), we built a homology model of the *C. elegans* ATP synthase. The ATP synthase dimer is too large to predict the structure as a single multimer, therefore we used AlphaFold (40) and AlphaFold multimer (41) to predict the structures of individual or small groups of subunits (Table S2, Fig. S6). Taking into account the fact that protein-protein interactions are likely important at the dimer interface, we predicted the dimer interface and peripheral stalk subunits both as individual subunits and as multimers. The peripheral stalk subunits were predicted successfully as a multimer, whereas the multimeric prediction for the dimer interface was poor. This could be explained by a limitation of AlphaFold multimer, which does not take stepwise assembly of complexes into account, instead assembling all proteins into a multimeric complex simultaneously (50). The result may also be attributable to the unique dimer interface in *C. elegans* compared to previously determined structures. We therefore used individual predictions to model the dimer interface (Fig. S6). The predicted *C. elegans* structures were then fitted sequentially into a scaffold provided by the *B. taurus* ATP synthase dimer [PDB 7AJB] (Fig. S7, Fig. 4A). The atomic model of *B. taurus* was chosen as a scaffold due to its closer relation to *C. elegans* (both being metazoans) and possessing an equivalent number of subunits in the c-ring. The *C. elegans* ATP synthase dimer model was then split into monomers and each was fitted sequentially into our sub-tomogram average dimer map (Fig. S7, Fig. 4B), improving the fit considerably (Fig. S8). The *C. elegans* homology model correlated well to the sub-tomogram averaging map (Fig. S9 and Table S3), providing us with a useful working model to allow a comparison of *S. cerevisiae* and *C. elegans* ATP synthase dimers (Fig. 4C).

**Figure 4.**
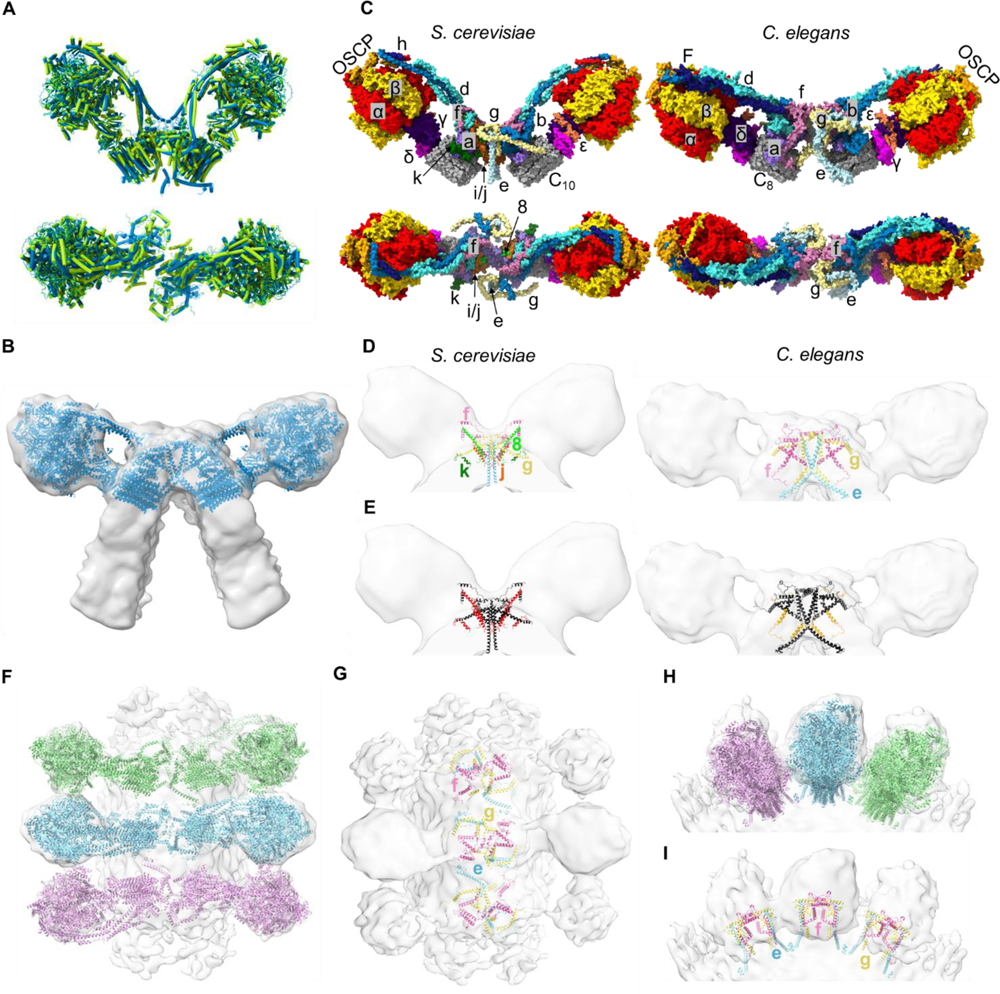
AlphaFold homology model of the *C. elegans* ATP synthase dimer. **(A)** AlphaFold predictions for *C. elegans* ATP synthase subunits (blue) overlaid with the atomic model of the bovine ATP synthase dimer ([PDB 7AJB] (29), green) that was used as a scaffold, using cylinder representation. Predicted models were fitted onto 7AJB using MatchMaker in ChimeraX. **(B)** Two monomers from the *C. elegans* ATP synthase homology model (helical representation) fitted into the sub-tomogram average of the *C. elegans* ATP synthase dimer. **(C)** Surface view of *S. cerevisiae* and *C. elegans* ATP synthase dimer models coloured by chain in side (top) and top-down (bottom) views. Subunits are annotated and shown as α, red; β, gold; γ, indigo; δ, magenta; ε, coral; c, grey; a, purple; b, blue; d, turquoise; F_6_, navy; OSCP, orange; e, pale blue; f, pink; g, yellow; j, brown; k, dark green; 8, lime. All subunits are labelled in the side views apart from subunit 8 which is buried. Only the dimer interface subunits are labelled in the top-down views. **(D)** Left, dimer interface subunits in the *S. cerevisiae* atomic model [6B8H] (29) coloured by chain and fitted into an *S. cerevisiae* sub-tomogram average [EMD-2161] (11). Right, dimer interface subunits in the *C. elegans* homology model coloured by chain fitted to the *C. elegans* sub-tomogram average. Subunits are annotated with the same colours as panel C. **(E)** As per (D), but with all subunits colored black, highlighting subunits missing in *C. elegans* relative to *S. cerevisiae* (j, k and 9) in red (left) and extensions in *C. elegans* subunits e, f and g relative to *S. cerevisiae* in orange (right). **(F)** Top-down view of the *C. elegans* ATP synthase dimer homology model fitted to the sub-tomogram average showing sequential dimer pairs in a row. **(G)** As per (F), but exclusively showing dimer interface subunits e, f and g coloured by chain as per panels C and D. **(H)** and **(I)** show the same interactions as in (F) and (G) respectively but viewed from the side of a dimer row.

Interestingly, the extra mass identified at the *C. elegans* dimer interface (Fig. 2B) appears to be filled by a rearrangement of the subunits f and g in the model (Fig. 4C, D & E). This agrees with the observation that these subunits show extensions relative to their yeast homologues (Fig. S10). In addition, extensions in the *C. elegans* peripheral stalk subunits (subunits b, d and F_6_) may also to contribute to the greater mass at the dimer interface compared to that observed in *S. cerevisiae* (Fig. S11). We cannot exclude the possibility that there are additional subunits as yet unidentified in *C. elegans* that may also contribute to the dimer interface. Finally, we fitted the *C. elegans* ATP synthase dimer model into a row of oligomeric dimer pairs along the curved edge of a crista (Fig. 4F, H). This reveals potential inter-dimer interactions mediated by subunit e (Fig. 4G, I and Fig. S12), in agreement with recent work demonstrating the key role that this subunit plays in oligomerisation and row formation (21).

## Discussion

Owing to the essential and universal role of the ATP synthase across eukaryotic species, it is remarkable that the dimeric interface can be so variable (10). Until now, the arrangement of ATP synthases in invertebrates was unknown, as was the correlation between dimer angle and whole mitochondrial morphology. In this work, a novel dimer angle for the ATP synthase from the nematode worm *C. elegans* was discovered. By comparing worm and yeast mitochondria, we correlated a wider ATP synthase dimer angle with flatter crista membrane morphology. Since dimer row formation is known to be instrumental in the formation of curved ridges in crista membranes (11,12,26), it is consistent that dimer angle influences the extent of membrane curvature.

The *C. elegans* ATP synthase dimer shows clear extra mass at the dimer interface when compared to other determined structures, which can be attributed to changes in subunit composition. Using sequence analysis, we detected extensions in 3 *C. elegans* dimer interface subunits (e, f and g), an extension in the peripheral stalk component subunit b, and a range of more subtle gaps and insertions in subunits d and F_6_. To investigate whether these could bulk out the width of the dimer interface, we built a homology model using AlphaFold (40) and AlphaFold multimer (41). A recently proposed alternative method employs the prediction of subcomponent structures using AlphaFold multimer based on known assembly intermediates (50). While conceptually advantageous for constructing a homology model of the ATP synthase dimer, only 50% of all high-resolution non-redundant complexes with 10-30 chains from the PDB were successfully assembled (50). Moreover, the efficacy of this approach has yet to be evaluated on protein complexes exceeding 30 chains. Our homology model of the *C. elegans* ATP synthase dimer thus allows us to hypothesise how alterations in the organisation of subunits could influence dimer architecture. The ATP synthase structure is relatively well conserved across species (51), but this conservation is weaker at the dimer interface and peripheral stalk (52). The extensions in *C. elegans* subunits e, f and g appear to result in the rearrangement of subunits at the dimer interface relative to *S. cerevisiae*. In addition, the extension in peripheral stalk component subunit b, and the changes to subunits d and F_6_, appear to bulk out the width of the dimer interface. Some dimer interface subunits present in *S. cerevisiae* (j, k and 8) are absent in *C. elegans.* Whilst it cannot be completely excluded that a yet unidentified subunit may substitute for subunit 8, we speculate that the absence of subunit 8 in worms (36) highlights an interesting evolutionary divergence. Subunit 8 is usually encoded by the mitochondrial genome, indicating that it is essential (38,39). Additionally, subunit 8 appears to have a key structural role in joining the dimer interface to the peripheral stalk (22,23,29,46). It is therefore plausible that the space vacated by the absent subunit 8 is either resolved by the re-arrangement of neighbouring subunits, or by substituting one of the extensions of the F_O_ subunits close by (b, d, e, f or g).

Mitochondria have evolved their highly convoluted crista membranes to increase their surface area (53), hence accommodating the maximum amount of respiratory chain complexes. This has made it possible for eukaryotic organisms to deal with higher energy demands than prokaryotes (53). A flatter crista (in *C. elegans*) compared to a wider one (in *S. cerevisiae*) could allow greater packing of respiratory chain complexes along the flat membrane surfaces (10), increasing the relative amount of proton pumping. It has been suggested that cristae serve as proton concentrators that facilitate a directed flow from the source (respiratory chain) to sink (ATP synthase) (9,10); protons have been proposed to preferably migrate from source to sink along membrane surfaces. If this were the case, then reducing the width of the crista space would reduce the solvent volume within which protons dissipate, facilitating the efficiency of ATP synthesis. Both these factors could allow *C. elegans* to maximise energy production in its soil-based habitat (54), where conditions range from near hypoxia to atmospheric (55,56). In summary, we propose that a wider ATP synthase dimer angle associated with flatter cristae may be paramount for capitalising on ATP production when a higher level of oxygen becomes available, and that a range of angles has evolved to meet the energetic needs of different organisms. Future studies geared towards investigating dimer subunit composition, angle and corresponding crista morphology across a range of species inhabiting different environments will be key in providing further support for this hypothesis. We demonstrate that the divergence in ATP synthase dimer architecture relative to yeast and mammalian systems makes *C. elegans* an ideal model system for further investigation of the role of dimer angle in mitochondrial physiology, health and disease.

## Materials and Methods

All standard reagents were purchased from Sigma-Aldrich (Burlington, USA).

### C. elegans and S. cerevisiae culture

The *C. elegans* N2 Bristol strain was maintained at 20°C on 60 mm Nematode Growth Medium (NGM) plates seeded with *E. coli* OP50. For large scale preparations, a semi-synchronised population of *C. elegans* (achieved by starving so that they entered the dauer stage) (57,58) were grown in a liquid suspension of *E. coli* NA22 in S-basal complete medium (59) at 20°C, shaking at 200 rpm for 3 days to achieve adults. For further details see (33). *S. cerevisiae* ‘Bakers’s yeast’ S288C derivative strains YPH499 were cultured at 19 −24°C in YPGal or YPG medium (1% w/v yeast extract, 2% w/v bactopeptone, 2% w/v galactose or 3% w/v glycerol) until OD 2-2.5 was reached. For further details see (60).

### Mitochondrial isolation

*C. elegans* and *S. cerevisiae* were both harvested from liquid cultures by low speed centrifugation. *C. elegans* preparation required an additional sucrose flotation step to remove debris. To soften the *C. elegans* cuticle, the pellets underwent collagenase treatment (1 U/ml collagenase, 100 mM Tris-HCl pH 7.4 and 1 mM CaCl_2_), whilst *S. cerevisiae* pellets underwent dithiothreitol (10 mM DTT, 100 mM Tris-SO4 pH 9.4) and zymolyase treatment (4.5mg/g zymolyase, 1.2 M sorbitol, 20 mM potassium phosphate, pH 7.4) to disrupt the cell wall. Pellets from both species were re-suspended in homogenisation buffers. For *C. elegans*, this was STEG/M (220 mM mannitol, 70 mM sucrose, 5 mM Tris-HCl pH 7.4 and 1 mM EGTA supplemented with 1 mM PMSF in methanol and 1% (w/v) fatty acid-free BSA). For *S. cerevisiae* the homogenization buffer contained 0.6 M sorbitol, 10 mM Tris–HCl pH 7.4, 1 mM PMSF, 0.2% (w/v) BSA, 2 mM magnesium acetate. The re-suspended *C. elegans* or *S. cerevisiae* samples were homogenised in a glass-Teflon Potter homogenisor to break open cells. Both samples were subsequently spun at low speed (750 – 3000 x g for 5-15 minutes) to remove cell debris and nuclei, and then at higher speed (12,000 x g for 15 minutes) to pellet mitochondria. Purified mitochondria were re-suspended in buffers that were optimised to maintain intact mitochondria: 220 mM mannitol, 70 mM sucrose, 5 mM Tris-HCl pH 7.4 and 1 mM EGTA for *C. elegans* or 250 mM sucrose, 2 mM magnesium acetate, 10 mM Mops, pH 7.2 for *S. cerevisiae*.

### Mitochondrial crista membrane isolation

Crista membranes used for the sub-tomogram averaging experiments were generated by successive freeze-thaw cycles of mitochondria at −80°C. To purify mitochondrial membranes from other cellular material, membrane extracts were incubated for 1h at 4°C with an anti-NDUFS3 primary antibody (ab14711; abcam) against the matrix arm of complex I from *C. elegans*, followed by a 3h incubation with an anti-mouse secondary conjugated to a quantum dot emitting at 625 nm (Q22085; Invitrogen). Crista membranes were separated from unbound antibodies and other cellular material on an Optiprep gradient with 10 layers (200 µl volume each) ranging from 0 to 27% v/v of iodixanol in STEG/M buffer, by centrifugation at 80,000 *× g* for 30 min at 4°C using a TLS-55 rotor (Beckman Coulter Inc., Miami, FL, USA). Crista membranes were identified and removed based on fluorescence under a UV lamp. Samples were then diluted in STEG/M buffer to wash out the iodixanol, and spun at 20,000 *x g* for 15 min at 4 °C to pellet the membranes. The enriched cristae were again re-suspended in STEG/M buffer.

### Electron cryo-tomography

Whole mitochondria or crista membranes were mixed 1:1 with 10 nm gold fiducials (Aurion, Wageningen, The Netherlands), applied to glow-discharged holey carbon EM grids (Quantifoil, Jena, Germany), and blotted for 5-6 seconds, followed by plunge-freezing in liquid ethane using a Vitrobot Mark IV (ThermoFisher, Massachusetts, USA) for *C. elegans*, or a home-made device for whole *S. cerevisiae* mitochondria. Pre-screening of *C. elegans* grids was carried out using an FEI Tecnai Spirit 120kV microscope (ThermoFisher), with a Oneview CCD Camera (Gatan, Pleasanton, USA). CryoET was performed using the same microscope for whole mitochondria, or using a 200 kV Talos Arctica (ThermoFisher) for crista membranes, equipped with a K2 direct electron detector camera and a GIF Quantum LS energy filter (Gatan). CryoET of whole *S. cerevisiae* mitochondria was performed using a 300 kV Titan Krios (ThermoFisher), K2 direct electron detector camera and a GIF Quantum LS energy filter (Gatan). Single tilt image series’ (±60, step size 1.5° −2°) were collected at -5 to -8 µm underfocus at nominal magnification of 21,000 x for whole mitochondria and 39,000 x for crista membranes, corresponding to 5.4 and 3.58 Å pixel sizes respectively for *C. elegans*, or 26,000 x for whole mitochondria from *S. cerevisiae*, corresponding to a 4.51 Å pixel size. The total dose per tomogram was ∼120 e^−^/Å^2^ for whole mitochondria, and ∼80 e^−^/Å^2^ for isolated cristae. Tomograms were aligned using the gold fiducials in IMOD (University of Colorado, United States) (61) and volumes reconstructed via weighted back-projection. Contrast was enhanced by nonlinear anisotropic diffusion (NAD) filtering (62), followed by manual segmentation, also in IMOD. ImageJ (63) was used to generate movies of segmentations generated in IMOD.

### Subtomogram averaging

3,234 *C. elegans* ATP synthase dimers were picked manually in IMOD, using NAD-filtered tomograms. Subvolumes containing the ATP synthase dimer were then extracted from tomograms that had not been NAD filtered. These sub-volumes were CTF corrected and imported into Relion 3.1 (64) using the approach and script described in (65). A reference-free initial model was generated using 3 x binned subvolumes and 2,481 dimers were selected by 2D classification for an unbinned refinement. Finally, 1,755 dimers were selected from a 3D classification of this refined model to enter a final round of refinement and post-processing, resulting in a 38.6 Å resolution map. Fig. S1 details the full workflow.

### Homology model generation

AlphaFold was used to predict five structural models of each ATP synthase subunit in *C. elegans* based on their mature protein sequence (40). Mature sequences were determined using MitoFates (66) or TargetP-2.0 (67) to predict mitochondrial targeting sequences. All ATP synthase subunits known to be present in *C. elegans* were included, excepting a putative homologue of subunit j, on account of its poor alignment with other homologues, and absence of any corresponding peptides in mass spectrometry analysis of the *C. elegans* dimer. The structures of peripheral stalk subunits b, d and F_6_ were predicted using AlphaFold multimer (41), as the individual predictions were unreliable. The models for each subunit with the highest average pLDDT score were fitted sequentially to a scaffold provided by the atomic model of the *S. cerevisiae* ATP synthase dimer [PDB 6BH8] in ChimeraX (68) using the Matchmaker tool. Where a subunit had more than one isoform, the version with the highest pLDDT score was used. In the case of subunit b, the isoform with the highest pLDDT score is also the only isoform expressed in somatic tissues (69). The resulting structure was divided into monomers, and fitted sequentially into the sub-tomogram average of the *C. elegans* ATP synthase dimer using the “fit in volume” tool in ChimeraX. The workflow is shown in Fig. S7. The resulting homology model was converted into an MRC map using the molmap command in ChimeraX (68). This map could then be fitted to the sub-tomogram average map of the *C. elegans* dimer for comparison (Fig. S19). The yeast monomeric atomic model [PDB 6CP6] (70) was used for additional analysis in Fig. S11.

### Mass spectrometry

The ATP synthase was purified from *C. elegans* mitochondria using a method described previously (71,72), and analysed by Nano-LC mass spectrometry. Briefly, isolated mitochondria were solubilised and mixed with a His-tagged inhibitor protein IF_1_. This suspension was applied to a Nickel column to capture inhibited ATP synthase. The fraction most enriched in ATP synthase subunits was taken for mass spectrometry analysis. Further details are given in Supporting Information.

## Data Availability

The sub-tomogram averaging maps generated in this study have been deposited in the Electron Microscopy Data Bank (EMDB) under accession code EMD-XXXX. The source image data have been deposited to the Electron Microscopy Public Image Archive (EMPIAR) under accession number [XXXX]. The Source Data accompanying Fig. 3B & E can be found in the accompanying Source Data file.

## Acknowledgments

We thank Rebekah White in the lab of Cameron Weadick for sharing equipment, resources and knowledge for ongoing nematode maintenance. We acknowledge Werner Kühlbrandt at the Max-Planck Institute of Biophysics, Frankfurt, Germany, where the *S. cerevisiae* data were collected. We thank Agnieszka Chacinska at IMol Polish Academy of Sciences, Warsaw, Poland, for supporting the *S. cerevisiae* based experiments. We acknowledge access and support of the GW4 Facility for High-Resolution Electron Cryo-Microscopy, funded by the Wellcome Trust (202904/Z/16/Z and 206181/Z/17/Z) and BBSRC (BB/R000484/1), and are grateful to Ufuk Borucu of the GW4 Regional Facility for High-Resolution Electron Cryo-Microscopy for help with screening and data collection. We thank Kate Heesom from the Bristol Proteomics Facility for collecting and analysing mass spectrometry data. EB was supported by the Biotechnology and Biological Sciences Research Council-funded South West Biosciences Doctoral Training Partnership [DTP2: BB/M009122/1] awarded to VG. MM was supported by a BBSRC responsive mode grant (BB/R008639/1) grant awarded to VG. PB was supported by the Foundation for Polish Science First TEAM Programme co-financed by the European Union under the European Regional Development Fund (POIR.04.04.00-00-3F36/17). BD received funding from the European Research Council (ERC) under the European Union’s Horizon 2020 research and innovation programme (grant agreement No 803894). This work was also funded by the Wellcome Trust (a Wellcome Investigator award (104632) to IC, which supported HF. The funders had no role in study design, data collection and interpretation, or the decision to submit the work for publication. For the purpose of Open Access, the authors have applied a CC BY public copyright license to any Author Accepted Manuscript version arising from this submission.

## Supporting Information Text

### Extended methods

### ATP synthase purification from *C. elegans* mitochondria

*C. elegans* ATP synthase was purified using a His-tagged IF_1_ as bait, following a scaled-down protocol designed for purification of bovine dimers (71,72). Residues 1-60 of the *C. elegans* F-ATPase inhibitor protein IF_1_ fused to a hexa-histindine tag (ceI1-60His), were overexpressed from a pRSFDuet plasmid in *E. coli* BL21 (DE3), and purified by affinity chromatography on a 5 mL Nickel-Sepharose column (Cytiva) attached to an ÄKTA purification system (Cytiva). Fractions enriched in IF_1_ were concentrated to ∼50 mg/mL with a VivaSpin concentrator (molecular weight cut-off 3 kDa; Sartorius).

*C. elegans* mitochondria were washed in a phosphate buffer (50 mM sodium hydrogen phosphate, 100 mM sucrose and 0.5 mM EDTA) and then centrifuged at 13,700 *x g* for 45 minutes at 4°C. This wash step was repeated twice to remove endogenous *C. elegans* IF_1_. Phosphate-washed mitochondria (∼16 mg) were solubilised for 30 minutes at 18°C at 7.65 mg/ml with digitonin (0.92% w/v) and DDM (0.76% w/v). The resulting extract was centrifuged at 24,000 *x g* for 20 minutes at 4°C, and ceI1-60His was added to the supernatant at 2.7 µg per 1 mg mitochondria to form ATPase:ceI1-60His complexes. A solution of 200 mM ATP, 200 mM MgSO_4_, and 400 mM Trizma (pH 8.0) was also added at 15 µl/ml before incubating for 15 minutes at 37°C, with further additions of this solution being added at 5 minute intervals. Precipitate was removed by centrifugation at 24,000 *x g* for 10 minutes at 4°C. NaCl and imidazole were added to the clarified sample to reach final concentrations of 150 mM and 25 mM respectively. This final extract was applied to a 1 mL HisTrap FF Nickel Column (Cytiva) installed on an ÄKTA purification system (Cytiva) and equilibrated in a buffer containing 20 mM Tris, pH7.4, 150 mM NaCl, 2 mM ATP, 2 mM MgSO_4_, 10% (v/v) glycerol, 0.1% (w/v) glyco-diosgenin (GDN) and a 0.1 mg/mL phospholipid mix. The ATPase:ceI1-60His complexes were eluted from the column by addition of a linear gradient of imidazole up to 500 mM over 10 mL. 0.5mL fractions were collected and run on an SDS-PAGE gel to confirm which fractions contained the ATPase:I1-60His.

### Nano-LC Mass Spectrometry

The sample of ATP synthase was run on a 10% SDS-PAGE gel until the dye front had migrated approximately 1cm into the separating gel. The gel lane was then excised as a single slice and subjected to in-gel tryptic digestion using a DigestPro automated digestion unit (Intavis Ltd.). The resulting peptides were fractionated using an Ultimate 3000 nano-LC system in line with an Orbitrap Fusion Lumos mass spectrometer (Thermo Scientific). Spectra were acquired with Xcalibur 3.0 software (Thermo Scientific).

The raw data files were processed and quantified using Proteome Discoverer software v2.1 (Thermo Scientific) and searched against the UniProt *Caenorhabditis elegans* database (downloaded October 2022; 26728 sequences) using the SEQUEST HT algorithm. Search criteria included oxidation of methionine (+15.995Da), acetylation of the protein N-terminus (+42.011Da) and methionine loss plus acetylation of the protein N-terminus (-89.03Da) as variable modifications and carbamidomethylation of cysteine (+57.021Da) as a fixed modification. Searches were performed with full tryptic digestion and a maximum of 2 missed cleavages were allowed. The reverse database search option was enabled and all data was filtered to satisfy false discovery rate (FDR) of 5%.

**Figure S1.**
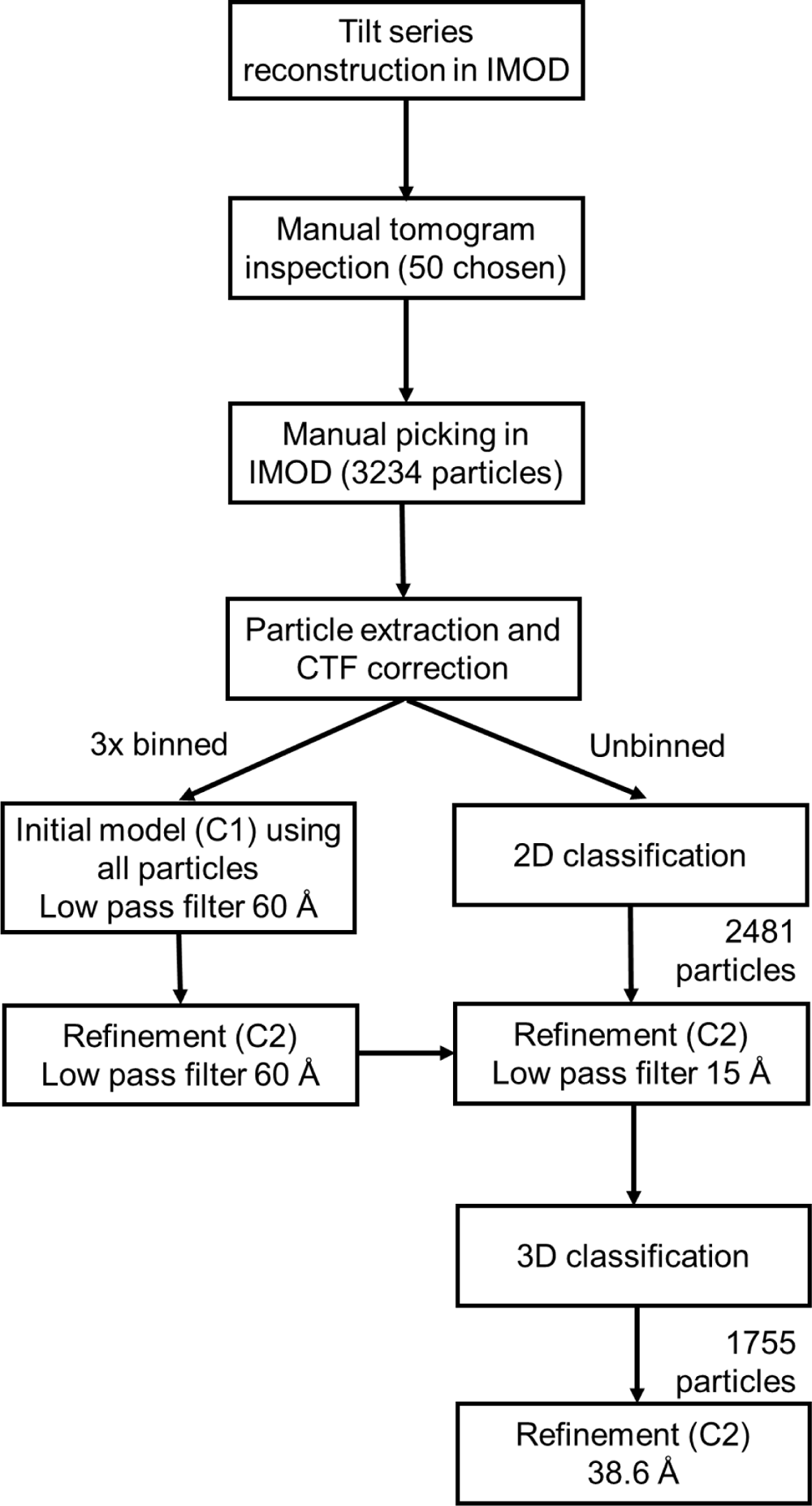
Flow chart of tomogram processing and sub-tomogram averaging using IMOD and Relion.

**Figure S2.**
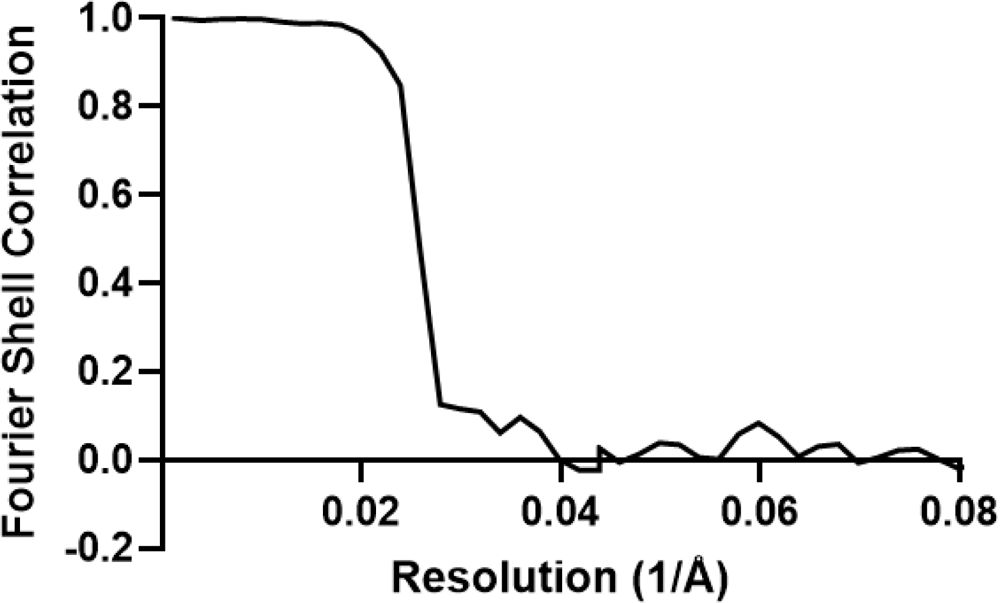
Fourier Shell Correlation (FSC) for the *C. elegans* ATP synthase sub-tomogram averaging map. The corrected FSC curve is an output from Relion 3.1 with a reported resolution of 38.6 Å.

**Figure S3.**
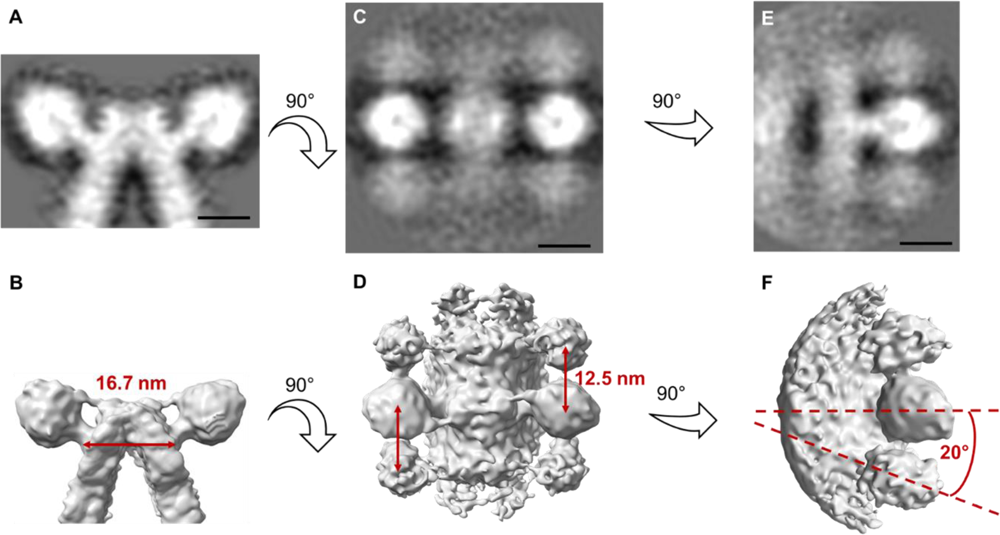
Inter-dimer distance and angle between consecutive dimer heads in oligomeric rows of *C. elegans* ATP synthase dimers. **(A)** 2D projection showing side view of a masked map of the *C. elegans* ATP Synthase dimer. **(B)** Side view shown in 3D, with distance between central stalks indicated. **(C)** 2D projection showing top-down view of an unmasked map of the *C. elegans* ATP synthase dimer. **(D)** Top-down view in 3D with inter-dimer distance indicated. **(E)** 2D projection showing side view (rotated 90° compared to A) of an unmasked map of the *C. elegans* ATP Synthase dimer. **(F)** Side view in 3D, with inter-dimer angle indicated. All indicated measurements were made in IMOD. Scale bars, 10 nm.

**Figure S4.**
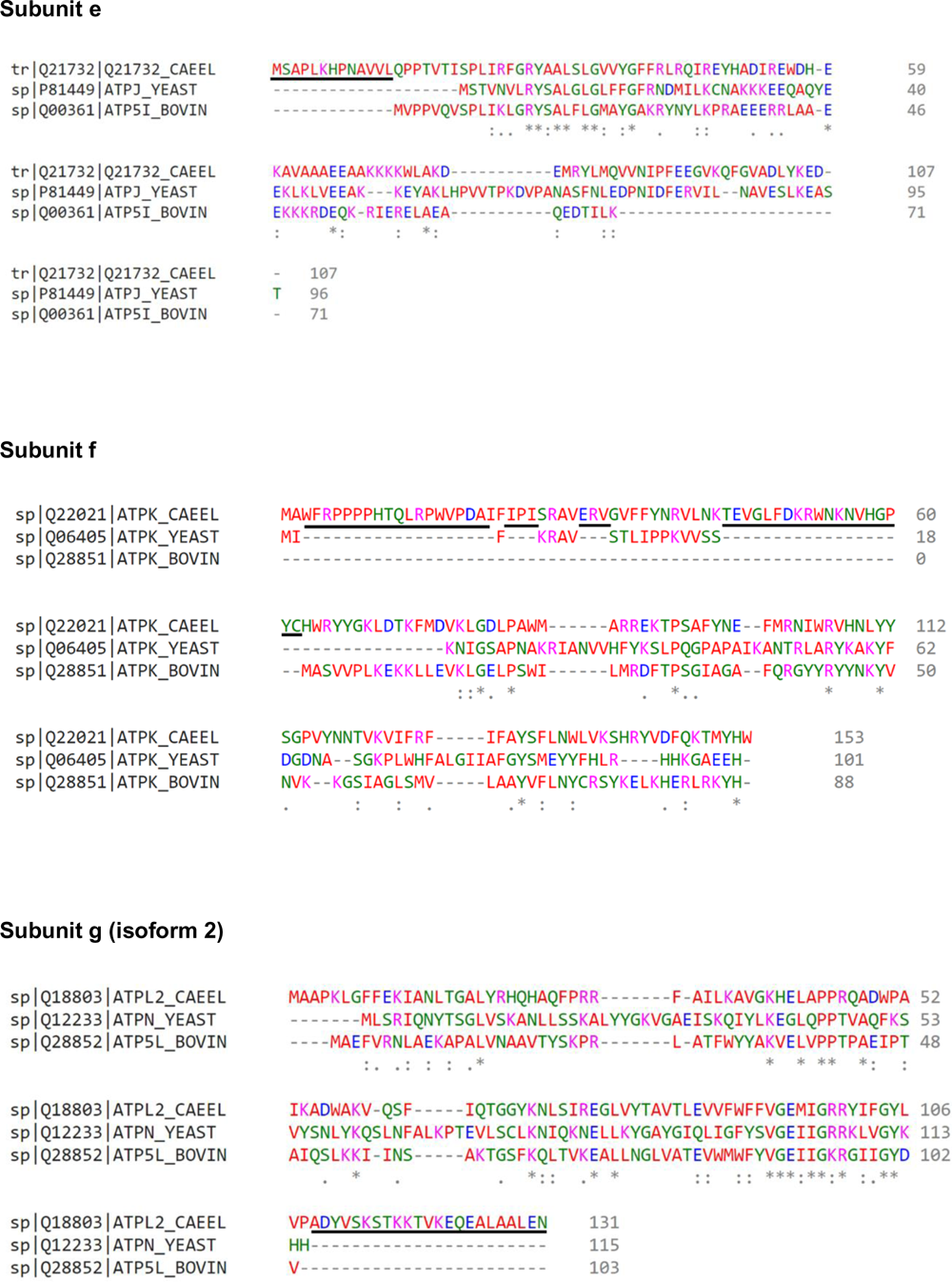
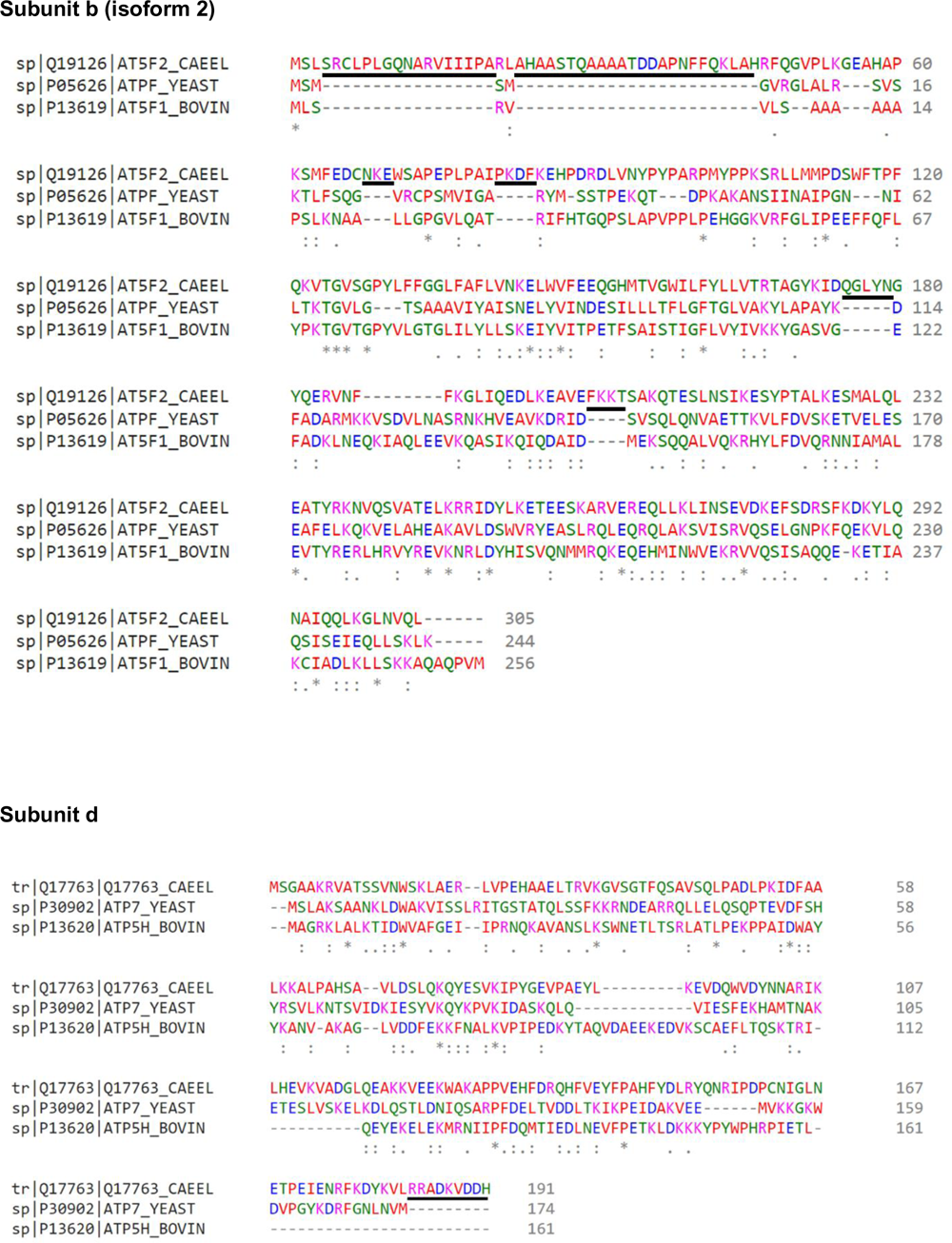
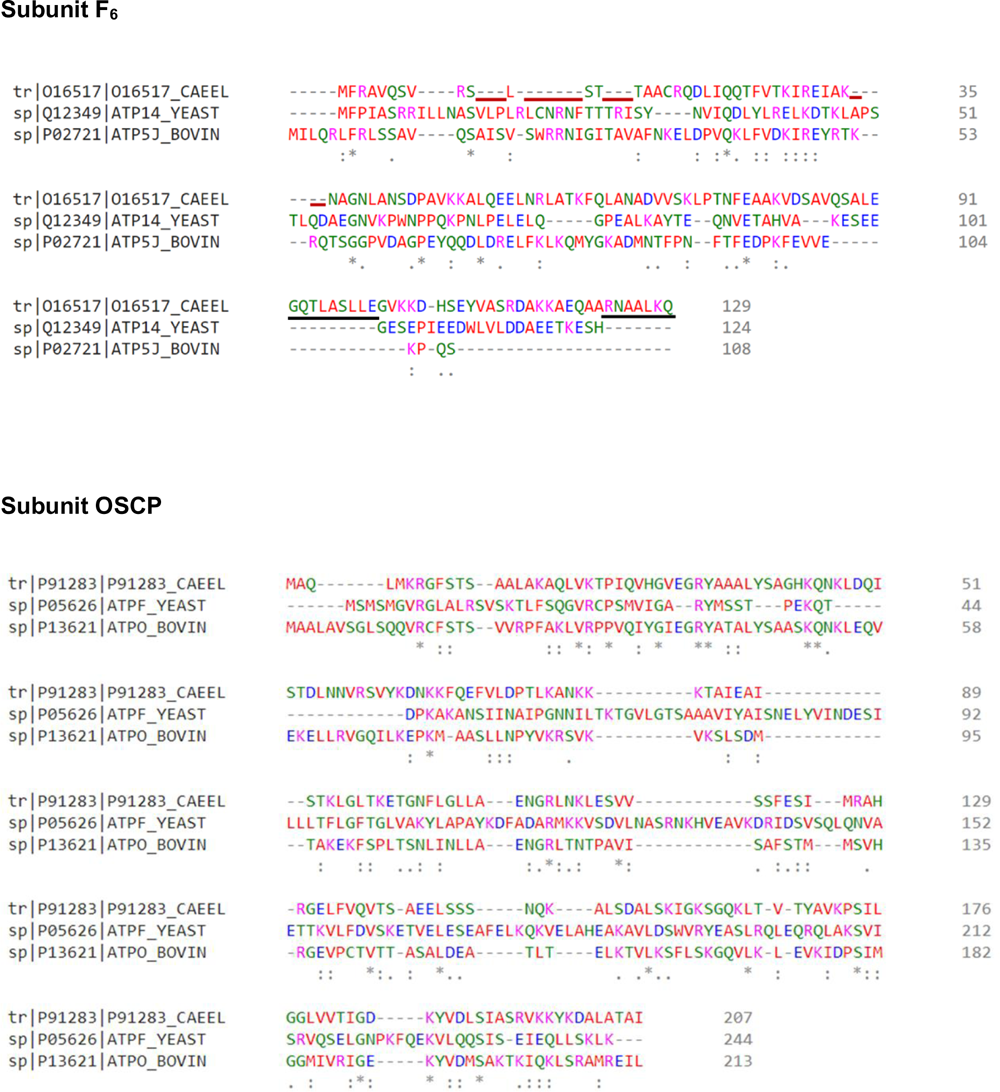
Multiple sequence alignment for dimer interface and peripheral stalk subunits. Comparisons were made between *C. elegans*, *S. cerevisiae* (Baker’s yeast strain ATCC 204508 / S288c) and *B. taurus* using Clustal Omega at EMBL-EBI (47–49). In the alignment output, an asterisk (*) indicates a perfect alignment, a colon (:) indicates a site belonging to a group exhibiting strong similarity, and full stop (.) indicates a site belonging to a group exhibiting weak similarity. Residues are coloured according to their biophysical properties. Small and hydrophobic residues are coloured red, acidic residues are coloured blue, basic residues are coloured magenta, and hydroxyl, sulfhydryl, amine and glycine residues are coloured green. Extensions in *C. elegans* subunits relative to both the *S. cerevisiae* and *B. taurus* homologues are underlined in black, deletions are underlined in maroon. Where subunits have multiple isomers, the isomer used in the homology model is used for alignment.

**Figure S5.**
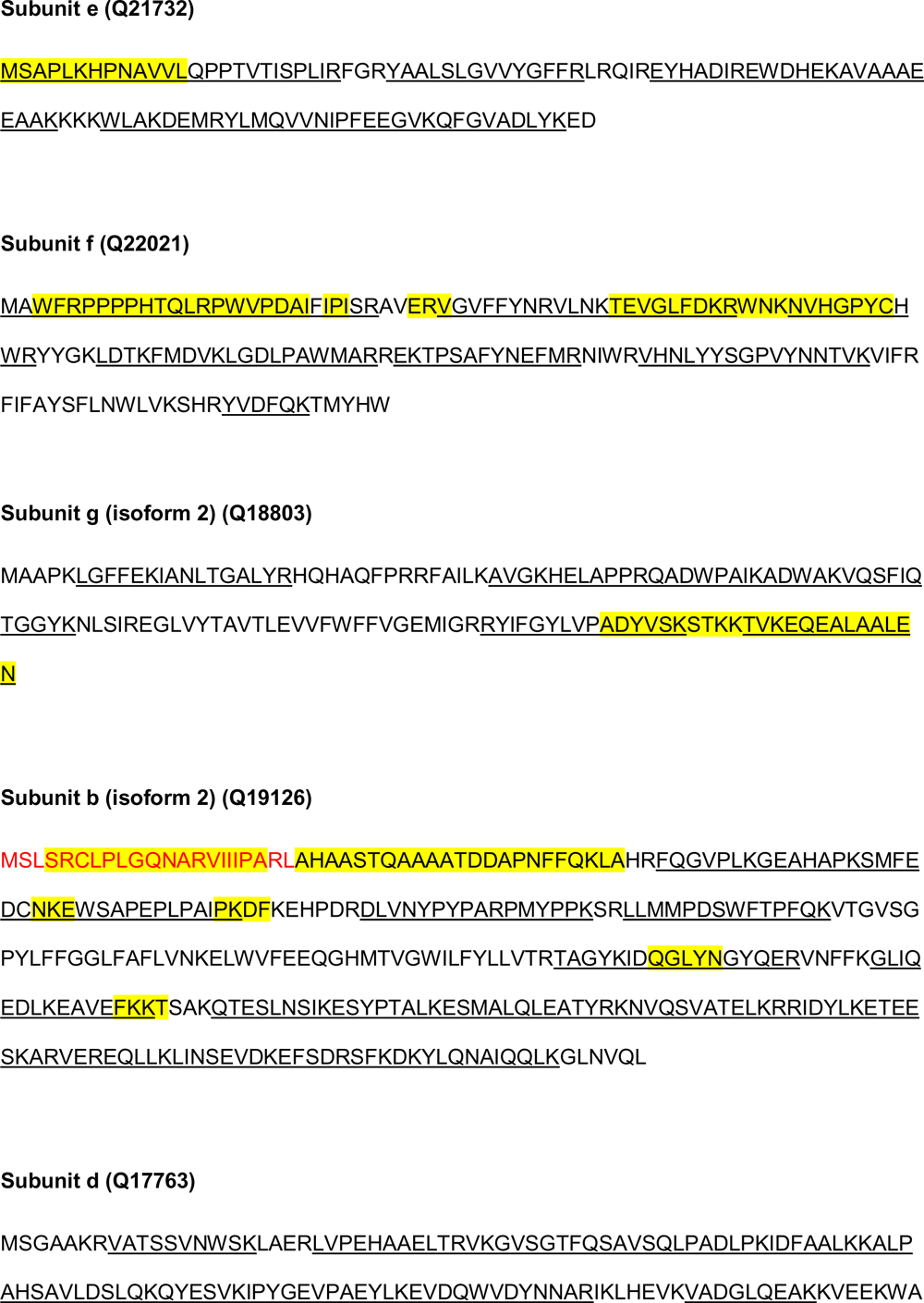
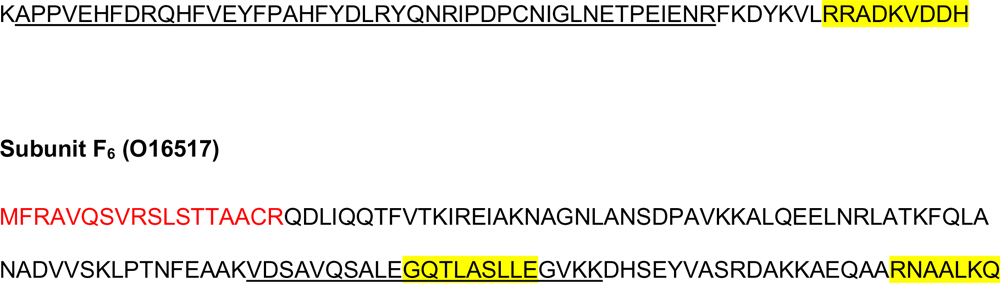
Mass spectrometry data for *C. elegans* ATP synthase subunits with significant extensions. The sequence for each subunit of interest is shown and identified with a Uniprot code. The predicted mitochondrial targeting sequences are coloured red. The *C. elegans* specific extensions (revealed in sequence alignments from Fig. S4) are highlighted in yellow. Peptides identified by mass spectrometry are underlined. Where subunits have multiple isomers, the isomer used in the homology model is shown.

**Figure S6.**
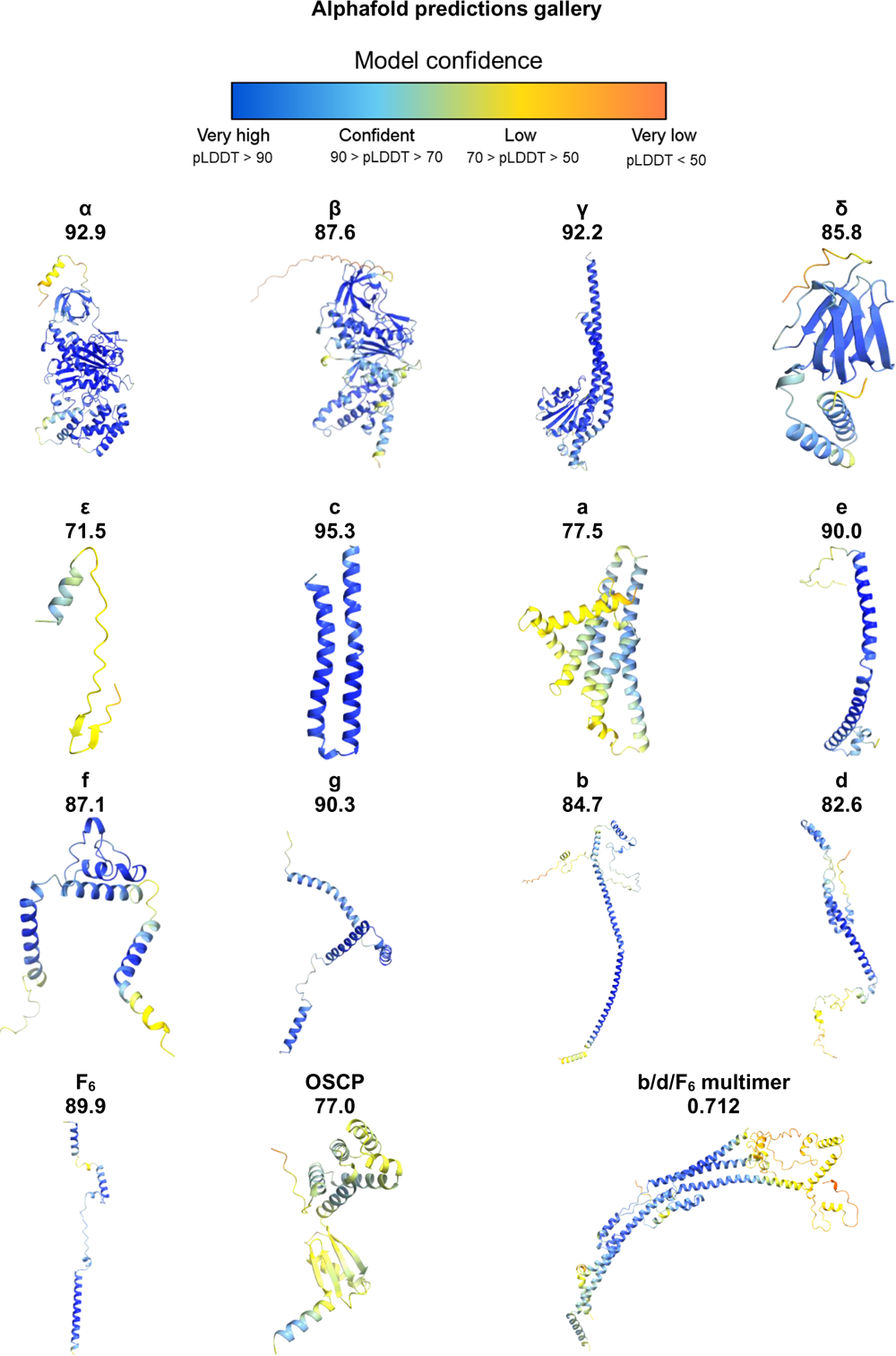
AlphaFold predictions gallery. AlphaFold predictions (40) for each *C. elegans* ATP synthase subunit, coloured by pLDDT score per residue. The pLDDT score is a per-residue measure of local confidence on a scale from 0 – 100. The structure of subunits b d and F_6_ were predicted as a multimer. The confidence measure for predictions made using AlphaFold multimer (41) is similar, but modified to score interactions between residues of different chains. It is calculated using a weighted combination of predicted-TM score (pTM) and interface predicted-TM score (ipTM), and has a scale from 0-1. The appropriate mean confidence score for each AlphaFold / multimer prediction is shown beneath each subunit name.

**Figure S7.**
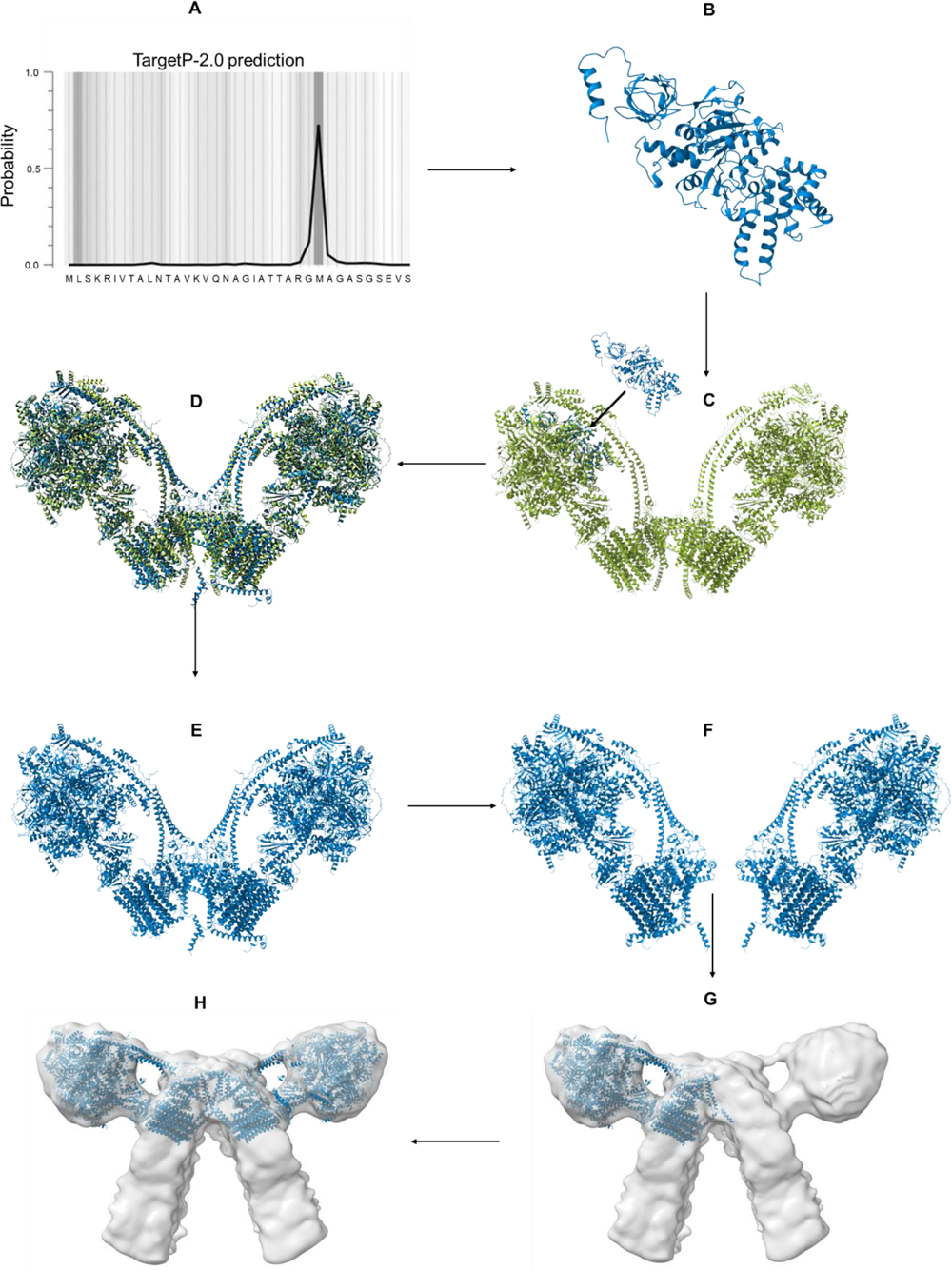
*C. elegans* ATP synthase homology model workflow. **(A)** MitoFates (66) or TargetP-2.0 (67) were used to predict the mitochondrial targeting sequence of individual proteins of the ATP synthase, so that the mature protein sequence could be identified. The example shown is the TargetP-2.0 prediction for subunit α. **(B)** AlphaFold was used to predict structures of all mature *C. elegans* ATP synthase subunits; again the example shown is the prediction for subunit α. **(C)** Predicted models were sequentially fitted into the *B. taurus* ATP synthase model [PDB 7AJB] (29) used as a scaffold using MatchMaker in ChimeraX. **(D)** The resulting homology model (blue) after all subunits have been fitted to the scaffold provided by 7AJB (green). **(E)** The homology model without the *B. taurus* scaffolding. **(F)** The *C. elegans* ATP synthase dimer was split into separate monomers. **(G)** The monomers were fitted sequentially into the sub-tomogram average of the *C. elegans* ATP synthase using matchmaker in ChimeraX to obtain the correct dimer angle. **(H)** The final homology model of the *C. elegans* ATP synthase dimer fitted into the sub-tomogram average.

**Figure S8.**
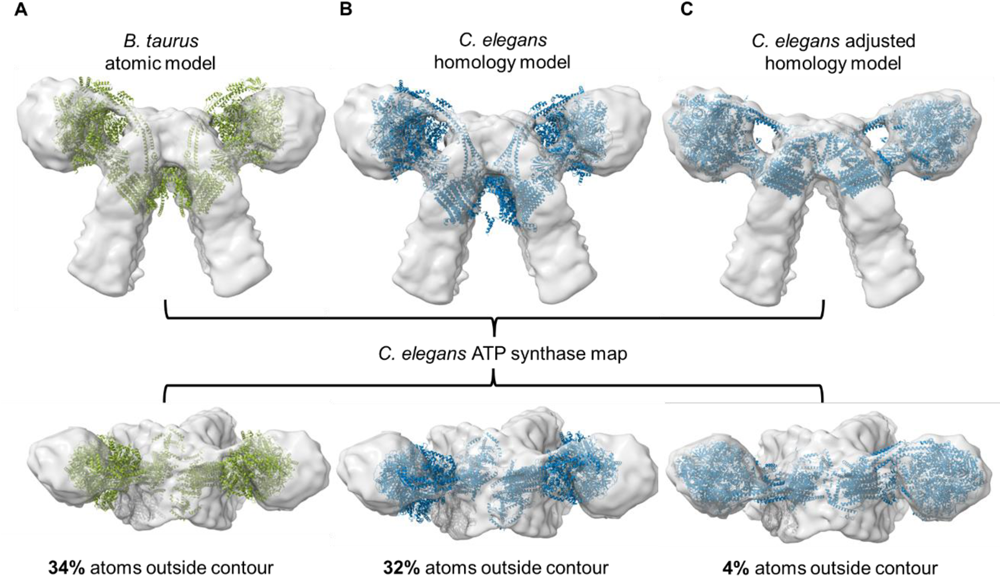
Comparison of different models fitted to the *C. elegans* ATP synthase dimer map. Different ATP synthase dimer models were fitted into the *C. elegans* ATP synthase *in situ* map. All models were fitted into the map at threshold 0.0429 in ChimeraX, and the percentage of atoms outside the contour is shown for each model. **(A)** The purified *B. taurus* ATP synthase dimer atomic model [PDB 7AJB] (29) used as a scaffold shows a poor fit, with 34% of atoms outside the contour. **(B)** The *C. elegans* ATP synthase dimer homology model following scaffolding to the *B. taurus* model also shows a poor fit, with 32% of atoms outside the contour. **(C)** Sequential fitting of monomers from the *C. elegans* homology model shows an improved fit, with only 4% of atoms outside the contour.

**Figure S9.**
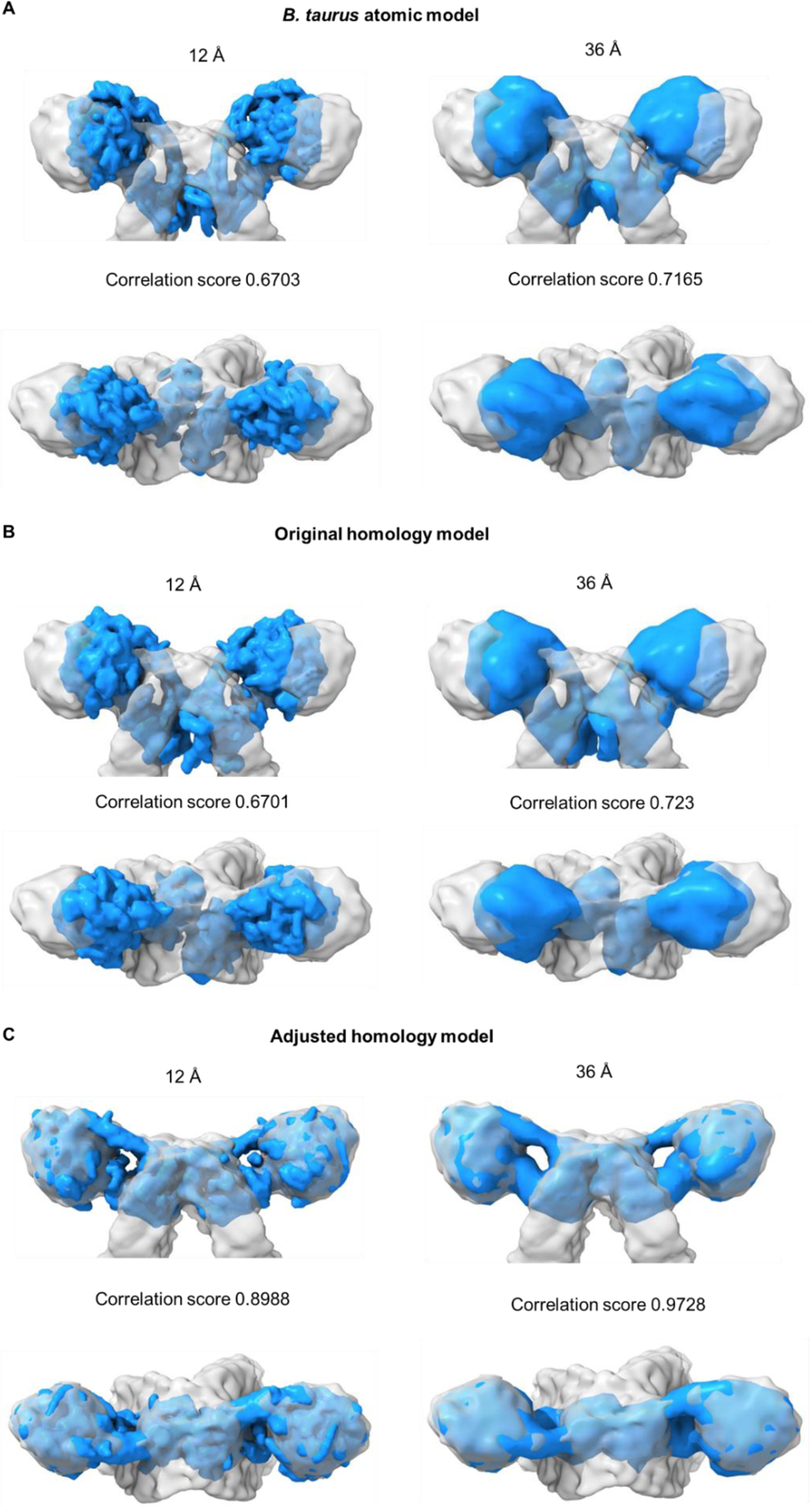
The *C. elegans* homology model fitted to the *C. elegans* ATP synthase dimer sub-tomogram averaging map. Using the molmap command in Chimera X (68), the PDB of the *C. elegans* homology model was converted into an MRC map at both 12 Å and 36 Å resolution. Converted molmap maps (blue) were then fitted to the sub-tomogram averaging map of the *C. elegans* dimer (grey) at equivalent threshold levels. Correlation scores between the homology model and sub-tomogram averaging maps are displayed. **(A)** Maps of the 7AJB *B. taurus* ATP synthase atomic model (29) used as a scaffold fitted to the sub-tomogram average for reference**. (B)** Maps of the *C. elegans* original homology model (without adjusting for dimer angle) fitted to the sub-tomogram average. **(C)** Maps of the dimer angle adjusted *C. elegans* homology model fitted to the sub-tomogram average.

**Figure S10.**
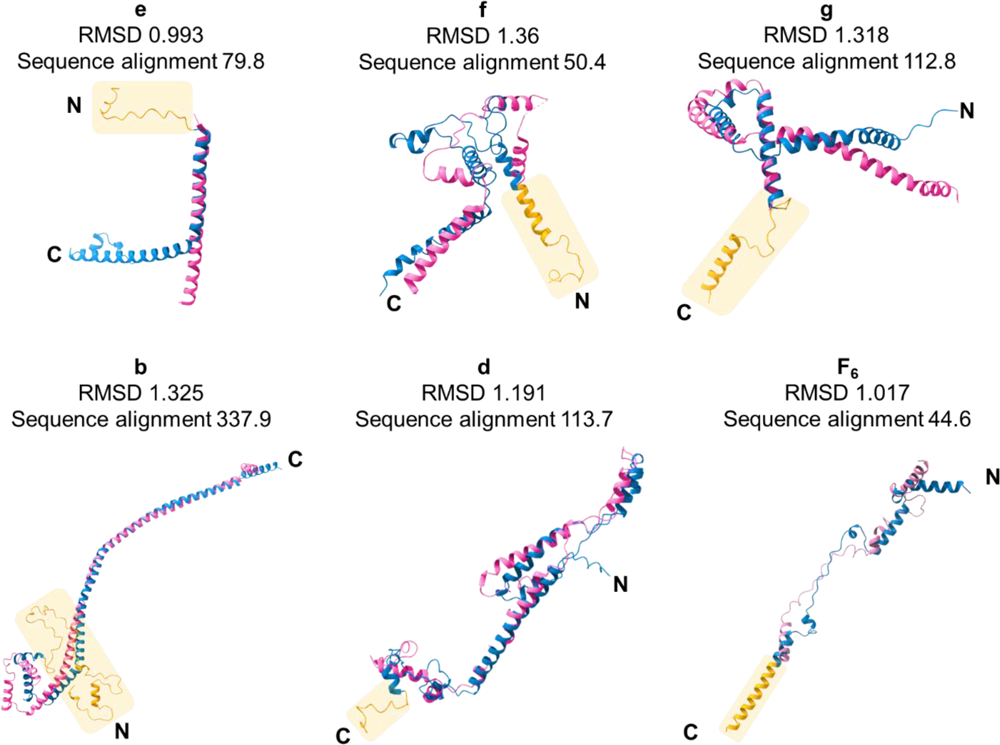
Overlays of individual subunits at the dimer interface and peripheral stalk, where there are extensions in *C. elegans* subunits compared with *S. cerevisiae*. *C. elegans* AlphaFold predictions (blue) are overlaid with their *S. cerevisiae* counterparts from the dimeric yeast ATP synthase atomic model ([PDB 6B8H], pink) (29). *C. elegans* subunit extensions are highlighted in orange. RMSD values (for pruned atom pairs) and sequence alignment scores output by ChimeraX when using the “fit to model” tool are shown for each overlay. Since the *S. cerevisiae* atomic model for the ATP synthase dimer [PDB 6B8H] does not contain complete density for subunit F_6_, the S. cerevisiae monomeric atomic model [PDB 6CP6] (70) was used to display a more complete *S. cerevisiae* chain for the overlay.

**Figure S11.**
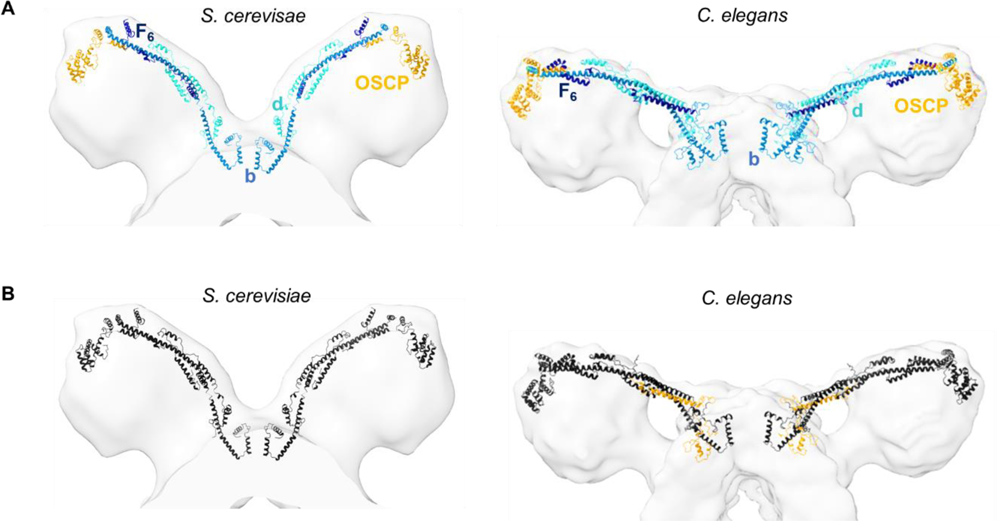
Comparison of peripheral stalk subunit arrangement in *S. cerevisiae* vs *C. elegans* ATP synthase dimers. **(A)** *S. cerevisiae* and *C. elegans* peripheral stalk subunits coloured by chain. Subunits are annotated and shown as b, blue; d, turquoise; F_6_, dark navy; and OSCP, orange. Left, peripheral stalk subunits b, d and OSCP in the 6B8H *S. cerevisiae* atomic model [PDB 6B8H] (29), and F_6_ from the monomeric atomic model [PDB 6CP6] (70), fitted to the *S. cerevisiae* sub-tomogram average [EMD-2161] (11). The chain for F_6_ was taken from 6CP6 (see Fig. S10B) as a large amount of density is missing from F_6_ in 6B8H (70). Right, *C. elegans* homology model fitted to the *C. elegans* sub-tomogram average. **(B)** As per (A), but with all subunits colored black, highlighting extensions in *C. elegans* subunits b, d and F_6_ relative to *S. cerevisiae* in orange.

**Figure S12.**
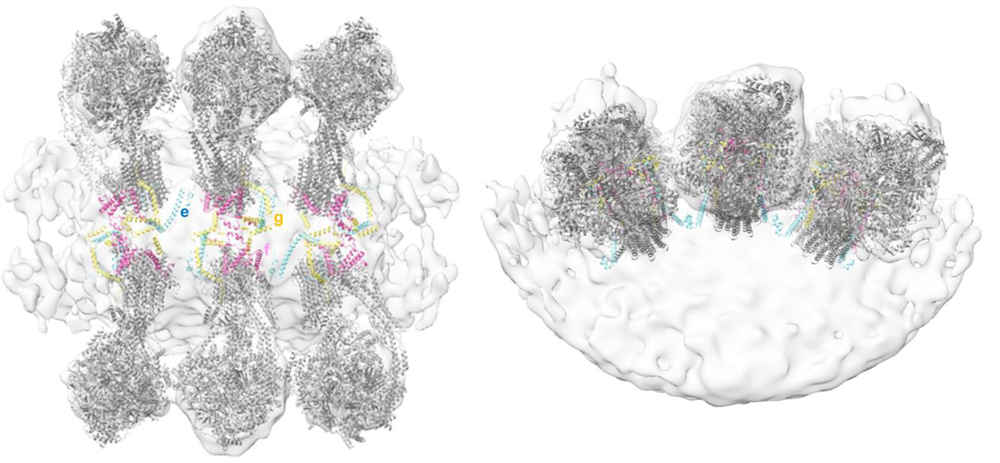
Inter-dimer interactions mediated by subunits e and g in *C. elegans* ATP synthase dimer rows. Top-down view (left) and side view (right) of the *C. elegans* ATP synthase homology model (grey) fitted to each dimer pair in the unmasked sub-tomogram average of the *C. elegans* dimer. Dimer interface subunits are colored (e, pale blue; f, pink; g, yellow) to highlight inter-dimer interactions mediated by subunits e and g.

**Table S1.** Nomenclature for homologues of ATP Synthase subunits.

| *C. elegans* | *S. cerevisiae* | *B. taurus* |
| --- | --- | --- |
| **F<sub>1</sub> head** |  |  |
| $\alpha$ | $\alpha$/Atp1 | $\alpha$ |
| $\beta$ | $\beta$/Atp2 | $\beta$ |
| **F<sub>0</sub> head** |  |  |
| $\gamma$ | $\gamma$/Atp3 | $\gamma$ |
| $\delta$ | $\delta$/Atp16 | $\delta$ |
| $\epsilon$ | $\epsilon$/Atp15 | $\epsilon$ |
| **Peripheral stalk** |  |  |
| b | b/Atp4 | b |
| d | d/Atp7 | d |
| F<sub>6</sub> | h/Atp14 | F<sub>6</sub> |
| OSCP | OSCP/Atp5 | OSCP |
| **F<sub>0</sub> motor** |  |  |
| a | a/Atp6 | a |
| c | c/Atp9 | c |
| **Type I dimer-specific subunits** |  |  |
| e | e/Atp21 | e |
| f | f/Atp17 | f |
| g | g/Atp20 | g |
| - | i/j/Atp18 | 6.8PL/ j |
| - | k/Atp19 | DAPIT/ k |
| - | 8/Atp8 | A6L/ATP8 |

Nomenclature for yeast and mammalian species are described as detailed by Song and Pfanner (6). In this work, we use primarily the mammalian nomenclature, which is also the standard used to describe *C. elegans* subunits. Exceptions to this are in our comparisons between *C. elegans* and *S. cerevisiae* dimers, where we use the *S. cerevisiae* naming system to describe subunits missing in worms.

**Table S2.** Metrics to assess confidence and fit of AlphaFold predicted structures.

| Subunit name | *C. elegans* Uniprot Accession Number | Mean pLDDT (or weighted pTM & ipTM)<sup>1</sup> | RMSD between pruned atom pairs<sup>2</sup> | RMSD across all atom pairs | Sequence alignment score<sup>3</sup> |
| --- | --- | --- | --- | --- | --- |
| α | Q9XXK1 | 92.9066444 | 0.586 | 1.009 | 2171.8 |
| β | P46561 | 87.5983733 | 1.161 | 3.202 | 2050.1 |
| γ | Q95XJ0 | 92.17635904 | 0.743 | 1.764 | 978.4 |
| δ | Q09544 | 85.82596715 | 0.901 | 0.941 | 368.7 |
| ε<sup>4</sup> | O16298 | 65.4116052 |  |  |  |
|  | P34539 | 71.51756013 | 0.631 | 6.924 | 39 |
| c | Q9BKS0 | 95.27668763 | 0.558 | 0.62 | 332.8 |
| e | Q21732 | 89.96603434 | 0.609 | 10.661 | 143.9 |
| f | Q22021 | 87.12810426 | 1.231 | 6.257 | 167.2 |
| g | Q18803 | 90.2590889 | 1.346 | 2.598 | 207.2 |
| a | P24888 | 77.54888203 | 1.073 | 4.582 | 322.9 |
| b | Q20053 | 84.43326886 |  |  |  |
|  | Q19126 | 84.74422485 | 1.076 | 8.068 | 441.4 |
| d | Q17763 | 82.60642993 | 1.175 | 4.902 | 296.7 |
| F<sub>6</sub> | O16517 | 89.9038886 | 0.79 | 6.717 | 76.2 |
| OSCP | P91283 | 76.95722866 | 1.09 | 1.557 | 517.7 |
|  | Q7JNG1 | 76.43462181 |  |  |  |
| b,d,F<sub>6</sub> multimer | Q19126, Q17763, O16517 | 0.712090029 | 1.076 | 8.068 | 441.4 |

<sup>1</sup> pLDDT scores are shown for subunits where structure was predicted individually, a weighted pTM and ipTM score is shown for a complex of subunits predicted using AlphaFold multimer. The pLDDT score is a per-residue measure of local confidence on a scale from 0 – 100. The predicted-TM score (pTM) and interface predicted-TM score (ipTM), and has a scale from 0-1.
<sup>2</sup> RMSD (Root Mean Square Deviation) is a measure of the similarity between two superimposed atomic coordinates, in this case for the predicted *C. elegans* subunits and the model of the *B. taurus* ATP synthase dimer.
<sup>3</sup> Sequence alignment score between *C. elegans* and *B. taurus*.
<sup>4</sup> Where a subunit has more than one isoform, the version with the highest pLDDT score was used to build the homology model. RMSD and sequence alignment scores are only shown for the selected protein. In the case of subunit b, the isoform with the highest pLDDT score is also the only isoform expressed in somatic tissues (69).

**Table S3.** Metrics to assess fit of atomic detail models to *C. elegans* ATP synthase dimer sub-tomogram averaging map.

|  | *B. taurus* atomic model [PDB 7AJB] | Original<sup>5</sup> *C. elegans* homology model | Adjusted<sup>6</sup> *C. elegans* homology model |
| --- | --- | --- | --- |
| PDB % atoms outside contour<sup>7</sup> | 34 | 32 | 4 |
| MRC map<sup>8</sup> correlation score | 0.7165 | 0.723 | 0.9728 |

<sup>5</sup> Homology model following scaffolding of AlphaFold predicted *C. elegans* subunits onto the *B. taurus* atomic model without adjusting for dimer angle.
<sup>6</sup> Homology model following fitting of dimer angle adjusted *C. elegans* ATP synthase monomers to the *C. elegans* ATP synthase sub-tomogram averaging map.
<sup>7</sup> This value is given by Chimera when fitting a PDB model to a map using the "fit in map" command.
<sup>8</sup> MRC map generated from PDB's using molmap command in ChimeraX (68). This metric shows level of correlation between molmap map and our sub-tomogram average at the same resolution (38.6 Å).

**Movie S1 (separate file).** Movie showing a 360° rotation about the y-axis of a single segmented *C. elegans* mitochondrion from the upper panel of Fig. 3A. An image sequence of 100 PNG files was collected in IMOD, and the sequence montaged into a10fps AVI file in Image J (63).

**Movie S2 (separate file).** Movie showing a 360° rotation about the y-axis of a single segmented *S. cerevisiae* mitochondrion from the lower panel of Fig. 3A. An image sequence of 100 PNG files was collected in IMOD, and the sequence montaged into a10fps AVI file in Image J (63).

